# Nanocall: An Open Source Basecaller for Oxford Nanopore Sequencing Data

**DOI:** 10.1101/046086

**Authors:** Matei David, L.J. Dursi, Delia Yao, Paul C. Boutros, Jared T. Simpson

## Abstract

**Motivation:** The highly portable Oxford Nanopore MinION sequencer has enabled new applications of genome sequencing directly in the field. However, the MinION currently relies on a cloud computing platform, Metrichor (metrichor.com), for translating locally generated sequencing data into basecalls.

**Results:** To allow offline and private analysis of MinION data, we created Nanocall. Nanocall is the first freely-available, open-source basecaller for Oxford Nanopore sequencing data and does not require an internet connection. On two *E.coli* and two human samples, with natural as well as PCR-amplified DNA, Nanocall reads have ~68% identity, directly comparable to Metrichor ”1D” data. Further, Nanocall is efficient, processing ~500Kbp of sequence per core hour, and fully parallelized. Using 8 cores, Nanocall could basecall a MinION sequencing run in real time. Metrichor provides the ability to integrate the ”1D” sequencing of template and complement strands of a single DNA molecule, and create a ”2D” read. Nanocall does not currently integrate this technology, and addition of this capability will be an important future development. In summary, Nanocall is the first open-source, freely available, off-line basecaller for Oxford Nanopore sequencing data.

**Availability:** Nanocall is available at github.com/mateidavid/nanocall, released under the MIT license.

**Contact:** `matei.david at oicr.on.ca`

## 1. INTRODUCTION

The MinION produced by Oxford Nanopore Technologies (ONT) is a highly portable, third-generation sequencing instrument, comparable in size to a cell phone. The small form factor makes the MinION particularly suitable for sequencing experiments performed in remote locations (Quick *et al*., 2016). However, the MinION relies on the cloud computing platform Metrichor (metrichor.com) for *basecalling, i.e*., translating the low-level, locally-generated sequencing data into DNA sequence reads. In a multi-site evaluation of the MinION using the SQK-MAP005 sequencing kit, Ip *et al*. (2015) obtained an average yield of ~115Mbp of ”2D” sequence data. This is encoded by Metrichor in more than 50Gb (basecalls and original events). One can easily envisage a setting in which the scarcity of internet access can limit the effectiveness of using a MinION for sequencing. Furthermore, the Metrichor source code is only available under a restrictive proprietary license, and we believe an open source basecaller would be valuable to the development community.

To address these limitations, we created and introduce here Nanocall, a basecaller for MinION sequencing data. Nanocall provides an offline alternative to Metrichor. In this sense, Nanocall has one important shortcoming compared to Metrichor, in that it performs strand-specific basecalls (”1D”, in Metrichor terminology), but it does not attempt to integrate the information from complementary strands (”2D”). We envisage three major use cases for Nanocall: 1) in situations where internet access is limited, such as remote sequencing; 2) as a rapid quality assessment check (*e.g*. that the correct sample was sequenced) prior to basecalling with Metrichor, allowing for real-time quality control; and 3) as an open source platform for testing new basecalling ideas and models.

ONT has recently announced the development of a new sequencing pore, R9. All data used in this paper is based on the currently available R7 sequencing pore using SQK-MAP006 sequencing kits.

## 2. BACKGROUND

### Sequencing overview

Informally, the MinION sequencer works as follows. First, DNA is sheared into fragments of 8-20 Kbp and adapters are ligated to either end of the fragments. The resulting DNA fragments pass through a protein embedded in a membrane via a nanometre-sized channel (this protein is the “nanopore”). A single strand of DNA passes through the pore; the optional use of a hairpin adapter at one end of the fragment allows the two strands of DNA to serially pass through nanopore, allowing two measurements of the fragment. In ONT terminology, the first strand going through the nanopore is the *template*, and the second is the *complement*. As a DNA strand passes through the pore it partially blocks the flow of electric current through the pore. The flow of current is sampled over time which is the observable output of the system. The central idea is that the single-stranded DNA product present in the nanopore affects the current in a way that is strong enough to enable decoding the electric signal data into a DNA sequence. This process, called *basecalling*, takes as input a list of current measurements, and produces as output a list of DNA bases most likely to have generated those currents.

### Segmentation

The first part of the decoding process is to segment the sampled current measurements into blocks. The nanopore current measurements are taken at reguiar time intervais, but the threading of the single-stranded DNA product through the nanopore is a stochastic process controlied by bioiogicai enzymes. The segmentation process takes as input the list of current measurements and produces a list of *events*, each consisting of: *start* `start`_*i*_, *length* `length`_*i*_, *mean* `mean`_*i*_, and *standard deviation* `stdv`_*i*_. Idealiy, each event corresponds to a different DNA context found inside the nanopore, and consecutive events correspond contexts differing by exactly one base. In practice, the segmentation process is noisy, so the event sequence will inevitably contain “stays” (consecutive events corresponding to the same context) and “skips” (consecutive events corresponding to contexts different by more than one base). The segmentation process is performed locally by the MinKNOW software running on the host computer.

### Pore Models

ONT provide *pore models* describing the events that are expected to be observed for various DNA contexts. For the SQK-MAP006 data used in this paper, the models use DNA sequences of length 6 (“6-mers”) as context, and for each 6-mer, they contain the *mean μ*_*k*_ and *standard deviation σ*_*k*_ of a Gaussian distribution modelling the *event mean*, and the *mean η_k_* and *standard deviation γ*_*k*_ of an Inverse Gaussian modelling the *event standard deviation*. These pore models are included in the basecalled `FAST5` files produced by Metrichor, and they depend only on the chemistry being used, not on the specific sample. There are currently three models in use, one for the template strand and two for the complement strand.

### Scaling Parameters

As a further complication, the current measurements have slightly different characteristics between nanopores, and between the times when they are taken by the same nanopore. To account for these variations, Metrichor uses a set of read-and strand-specific scaling parameters: `shift`, `scale`, `drift`, `var`, `scale`′, and `var`′. Using these parameters, if event *i* corresponds to context *k*, then:

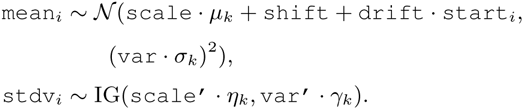

### Basecalling

The core of the decoding process is the basecalling step, performed in the cloud by Metrichor, which infers the DNA sequence most likely to have produced the observed event sequence. Metrichor currently uses a Hidden Markov Model (HMM), where the hidden state corresponds to the DNA context present in the nanopore, and where the pore models are used to compute emission probabilities. HMMs have been used before to model Oxford Nanopore data (Timp *et al*., 2012; Schreiber and Karplus, 2015; Loman *et al*., 2015; Szalay and Golovchenko, 2015). We describe our approach to basecalling in the following section.

## 3. METHODS

Our motivation behind Nanocall is to offer an offline alternative to Metrichor, and since the initial signal segmentation step is performed locally, we do not seek to replace it. As such, the input to Nanocall consists of a set of segmented event sequences, stored in ONT-specific `FAST5` files.

Nanocali can also be given a set of state transition probabilities and a set of pore models, but these are optional, and if they are not specified, sensible defaults are used. Nanocall processes each input file separately as follows. It begins by splitting the template and complementary strands into separate event sequences when a hairpin is found. Next, it estimates the pore model scaling parameters. Optionally, Nanocall can perform several rounds of training to update the scaling parameters using the Expectation Maximization algorithm, and also to update the state transition parameters using the standard Baum-Welch algorithm (Baum, 1972). Finally, Nanocall performs standard Viterbi decoding of the path through the hidden states, where the state is the 6-mer in the nanopore. The details of the individual steps are given below. For a reference on standard HMM algorithms, see Durbin *et al*. (1998).

### State Transitions

The state transitions are prior probabilities of moving from one state to another state. The default state transitions are computed based on two parameters: the “stay” probability Pstay and the “skip” probability *p*_skip_. The former, *p*_stay_, is the probability that two consecutive events are emitted from the same context/state. This corresponds to a segmentation error where an erroneous event break was introduced. The latter, *p*_skip_, is the probability that two consecutive events are emitted from states that differ by more than one kmer shift. This corresponds to either a segmentation error or a sequencing error (*i.e*., the DNA moved too quickly through the pore to register a detectable event) where one or more events were lost. A slight complication is that, by increasing the number of skips, there is always more than one way of going from any one state to any other. For example, `ACGTGT` can be followed by `GTGTAC` using either one or three skips. Our computation of the state transitions takes this into account:

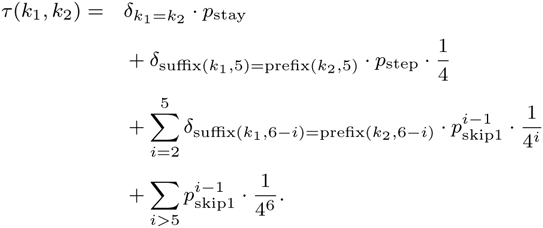

Here: *δ* is the standard indicator function; prefix(*k*, *i*) / suffix(*k*, *i*) are the prefix/suffix of *k* of length *i*; *p*_step_:= (1 – *p*_stay_ – *p*_skip_); and *p*_skip1_ = *p*_skip_/ (1 % *p*_skip_) corresponds to the probability of exactly one skip.

To speed up computation, for both Forward-Backward (used during training) and Viterbi (used during basecalling) algorithms, we disregard transitions between states corresponding to more than one skip. Thus, each state has at most 21 neighbours: itself, 4 at distance 1, and 16 at distance 2.

By default, Nanocall uses *p*_stay_ = .1 and *p*_skip_ = .3. When transition parameter training is enabled (see Supplement), the values of *p*_stay_ and *p*_skip_ are updated individually for each strand of each read, using the Baum-Welch algorithm.

### Pore Models

Nanocall is designed to work by default with the (6-mer based) pore models provided by Metrichor. These are three pore models, one for the template strand and two for the complement strand. Emission probabilities are calculated by multiplying the probability density from the Gaussian and Inverse Gaussian distributions that model the event mean and event standard deviation, respectively. This is done after the models are scaled (see Scaling below.) Nanocall can also run with user-provided pore models, though changing the kmer size requires recompilation.

### Strand Separation

The segmented list of events available in pre-Metrichor `FAST5` files does not always contain markers delimiting the template and complement strands. To deal with this, Nanocall implements a simple heuristic for separating the strands. The core idea here is that the “hairpin” adapter connecting the strands (in the single-stranded DNA product being threaded through the nanopore) contains abasic DNA, that is, DNA backbone lacking DNA bases. This abasic DNA generates a specific signal in the event sequence.

In the first step, we use a heuristic to detect the *abasic current level*. In general, the currents corresponding to regular DNA are in the range 50-90 pA, and those corresponding to abasic DNA are higher than 100 pA. However, the exact levels are affected by the `shift` and `scale` parameters, which are not known *a priori*. For that reason, Nanocall uses a heuristic to estimate abasic current level. Specifically we take the current level larger than 99% of all event levels in the read, and add 5pA.

In the second step, Nanocall looks for islands of 5 or more consecutive abasic current measurements. Then, islands within 50bp of each other are merged. Next, Nanocall selects the island closest to the middle of the event sequence. If this island is within the middle third of the entire event sequence it is used to separate the events corresponding to the two strands. If on the other hand this island is outside of the middle third of the event sequence, Nanocall gives up trying to separate the strands and attempts to basecall the entire event sequence as a template strand.

### Scaling

Pore model scaling parameter estimation is a crucial part of the basecalling process. Since accurate scaling dominates the running time, Nanocall presents the user with several scaling options.

In all cases, Nanocall initializes the scaling parameters using a crude “Method of Moments” approach, that fits the mean and standard deviation of (a subset of) the event sequence to the mean and standard deviation of the pore model being scaled. This approach relies on the assumption that the states producing the observed events are sampled uniformly at random, which is clearly not the case. However, for event sequences that are long enough, this method provides a reasonable base setting. This method only estimates 2 of the 6 scaling parameters discussed in the Background section: `shift` and `scale`. The remaining 4 are left with default values (0 for additive terms, 1 for multiplicative factors). The MoM scaling is the fastest, but also the least accurate. Since the other available scaling options dominate the total runtime, Nanocall can be instructed to use only MoM scaling, which will result in the fastest operation (labelled `fast`).

Beyond MoM, Nanocall offers the option to train the pore model scaling parameters by performing several rounds of an Expectation Maximization (EM) algorithm. The training stops either when the maximum allowed number of rounds is reached (by default 10 for single-strand scaling/20 for double-strand scaling, see below), or when the improvement in likelihood drops below a certain threshold (by default, a multiplicative factor, e). Each training round proceeds as follows. First, the “E” step consists of running the Forward-Backward algorithm to compute state posterior probabilities on 2 subsets of the event sequence, extracted from the start and end of each strand. Next, the “M” step updates all 6 scaling parameters with values that maximize the likelihood of the observed emissions given the probabilistic state assignments. More details about this update process are given in the Supplement.

Nanocall provides the user with the option of performing either single‐ or double‐ strand scaling. With single-strand scaling, the 2 strands of each read are processed independently of each other. The motivation for doublestrand scaling is to avoid potential overfitting of the two strands separately by constraining the scaling parameters to be the same across the read.

### Viterbi Decoding

After any optional training, Nanocall runs the Viterbi decoding algorithm to compute the most likely state sequence to have generated the observed event sequence. Then, the final base sequence is constructed by iteratively adding the minimum number of bases required to transition between consecutive states. Thus, for instance, if 2 consecutive states are `ACTCTC` and `CTCTCA`, the base sequence produced will be `ACTCTCA`, not ACTCTCTCA. In particular the basecalled sequence will never contain a homopolymer longer than 6bp as a result of this heuristic.

When several pore models are applicable to the same strand, as is the case with the default models and the complement strand, Nanocall will select which model to use during training if (training is performed, and) the Forward-Backward computed likelihood using one model is better than all other applicable models by a multiplicative factor of e^20^. If a model is not selected during training, the read strand is basecalled using all applicable models. Then, the model used for output will be the one with a higher joint probability of the most probable path through the HMM and the observations, found in the last cell of the Viterbi matrix Durbin *et al*. (1998).

### Evaluations

In our evaluations, we use BWA MEM with options ‘-x ont2d‘ for mapping ONT reads to the reference (Li, 2013). A 1D read (Metrichor or Nanocall) is considered *mapped* if its BWA MEM mapping overlaps the mapping of the corresponding Metrichor 2D read. Otherwise, the 1D read is *mismapped* (i.e., it is not mapped, or mapped elsewhere). The *identity* of a mapped read is defined as its mapping edit distance divided by the read length. The latter includes any clipping, so the basecaller is penalized for producing incorrect read ends. In our results, we only measure identity for reads where all 4 1D reads (2× Nanocall, 2× Metrichor 1D) are mapped.

### Source Code

Nanocall is written in C%%11 and its source code is available at github.com/mateidavid/nanocall, along with instructions for how to build it either in a standard UNIX environment, or as a Docker container. The analysis scripts used to generate the data presented in this paper are available at github.com/mateidavid/nanocall-analysis.

### Data Availability

The datasets used in this paper are available in the European Nucleotide Archive under accession numbers ERR1147227 (*E.coli* native), ERR1147229 (*E.coli* PCR), TBD (human native), and TBD (human PCR). The *E.coli* datasets were desribed by Quick *et al*. (2014).

## 4. RESULTS

We ran Nanocall on four ONT datasets consisting of *E.coli*, PCR-amplified *E.coli*, human, and PCR-amplihed human DNA. For each dataset, we used the first 10,000 reads labelled as “passing” by Metrichor, and ran Nanocall using 5 different option sets: no training (`fast`), and single/double strand scaling with/without transition parameter training ({`1/2`}{`ss/ss-no_tt`}). The main results are given in Table 1. One important conclusion supported by these runs is that the `fast` mode, with no training at all, seems very adequate for *E.coli* data, but much less so for human data. The difference between the two is the size and complexity of the genome, with *E.coli* reads being much easier to map. Another conclusion we were able to draw is that double strand scaling (`2ss`) performs better than single strand scaling (`1ss`). While the effect is minimal on *E.coli* data, it becomes quite pronounced on human data, where single strand scaling leads to an additional 4–6% reads being mismapped. When it comes to transition parameters, surprisingly, no training performs slightly better than training on human data: 2% on human native data, < 1% on human PCR data. However, not training transition parameters makes the performance of a run much more dependent on the default transition parameters. For this reason, and in spite of the slight increase in mismapping rate, we decided on using double strand scaling with transition parameter training (`2ss`) as the Nanocall default.

**Table 1.**
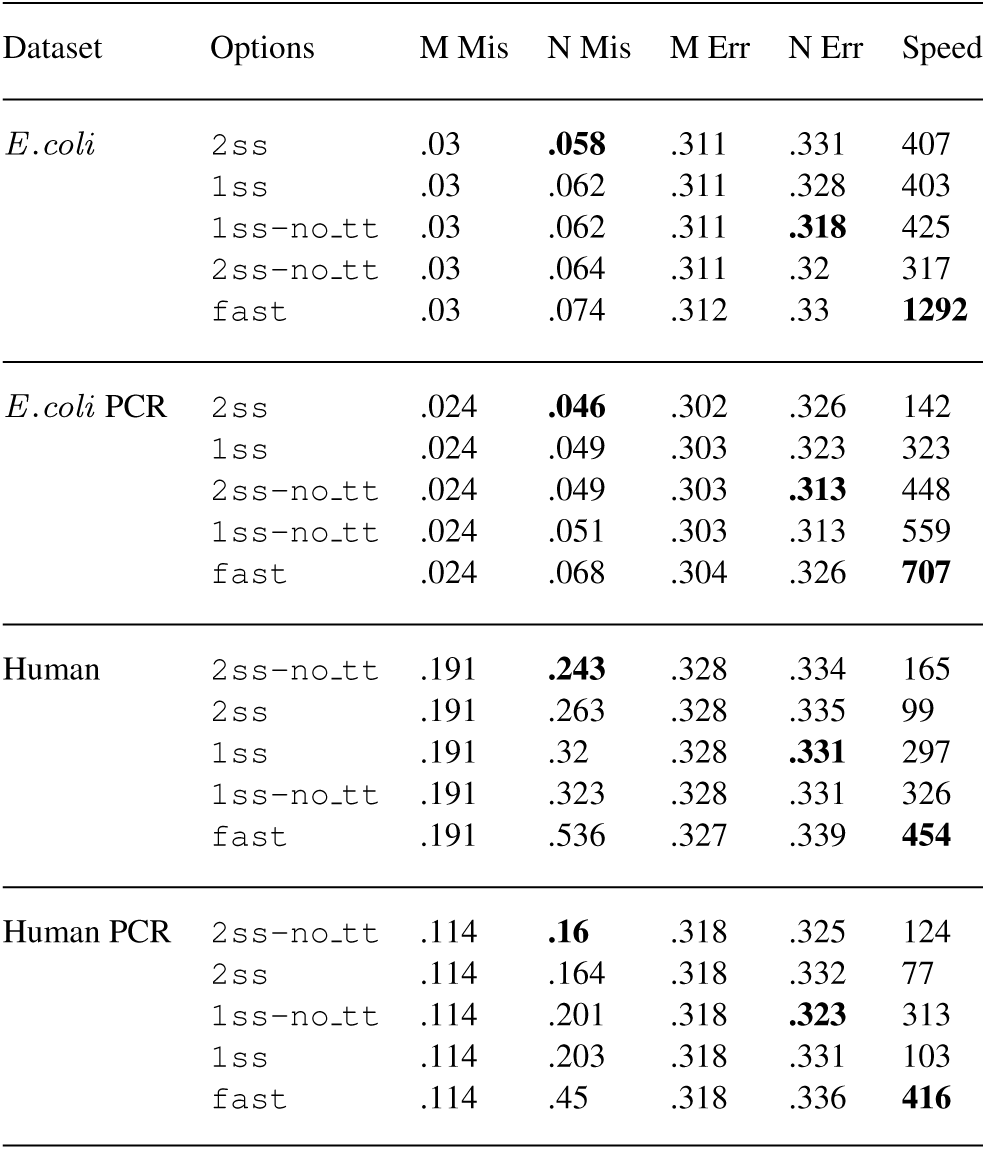
Performance of Nanocall. Options: `fast`: no training; {`1/2`}{`ss/ss-no_tt`}: single/double strand scaling, with/without transition parameter training. M/N Mis: fraction of Metrichor 1D/Nanocall reads where the mapping of at least one strand does not overlap the mapping of the Metrichor 2D read. M/N Err: Mapping edit distance divided by read length, for reads where all 5 mappings (1× Metrichor 2D, 2× Metrichor 1D, 2× Nanocall) are overlapping. Speed: Kbp per core-hour.

| Dataset | Options | M Mis | N Mis | M Err | N Err | Speed |
| --- | --- | --- | --- | --- | --- | --- |
| *E.coli* | 2ss | .03 | **.058** | .311 | .331 | 407 |
|  | 1ss | .03 | .062 | .311 | .328 | 403 |
|  | 1ss-no_tt | .03 | .062 | .311 | **.318** | 425 |
|  | 2ss-no_tt | .03 | .064 | .311 | .32 | 317 |
|  | fast | .03 | .074 | .312 | .33 | **1292** |
| *E.coli* PCR | 2ss | .024 | **.046** | .302 | .326 | 142 |
|  | 1ss | .024 | .049 | .303 | .323 | 323 |
|  | 2ss-no_tt | .024 | .049 | .303 | **.313** | 448 |
|  | 1ss-no_tt | .024 | .051 | .303 | .313 | 559 |
|  | fast | .024 | .068 | .304 | .326 | **707** |
| Human | 2ss-no_tt | .191 | **.243** | .328 | .334 | 165 |
|  | 2ss | .191 | .263 | .328 | .335 | 99 |
|  | 1ss | .191 | .32 | .328 | **.331** | 297 |
|  | 1ss-no_tt | .191 | .323 | .328 | .331 | 326 |
|  | fast | .191 | .536 | .327 | .339 | **454** |
| Human PCR | 2ss-no_tt | .114 | **.16** | .318 | .325 | 124 |
|  | 2ss | .114 | .164 | .318 | .332 | 77 |
|  | 1ss-no_tt | .114 | .201 | .318 | **.323** | 313 |
|  | 1ss | .114 | .203 | .318 | .331 | 103 |
|  | fast | .114 | .45 | .318 | .336 | **416** |

Overall, Table 1 shows that the reads produced by Nanocall (`2ss`) are directly comparable to Metrichor 1D reads: Nanocall increases the mismapping rate by an additional 3% for *E.coli* data, and 6% on human data. Furthermore, Nanocall and Metrichor 1D reads have largely similar percent identity at 68%. The similarity between Nanocall and Metrichor 1D reads is further demonstrated in Figures 1-9, where we compare Nanocall reads with Metrichor 1D (same strand) and 2D reads, on several metrics: scaling parameters `scale` and `shift`, identify, read length, and fraction aligned. The only concern in here could be a subpopulation of reads in Figure 1 on which Metrichor `scale` is notably smaller than Nanocall `scale` (slightly more pronounced in human data). This could be partly explained by the fact that Nanocall performs double strand scaling, while (we believe) Metrichor performs single strand scaling.

**Fig. 1.**
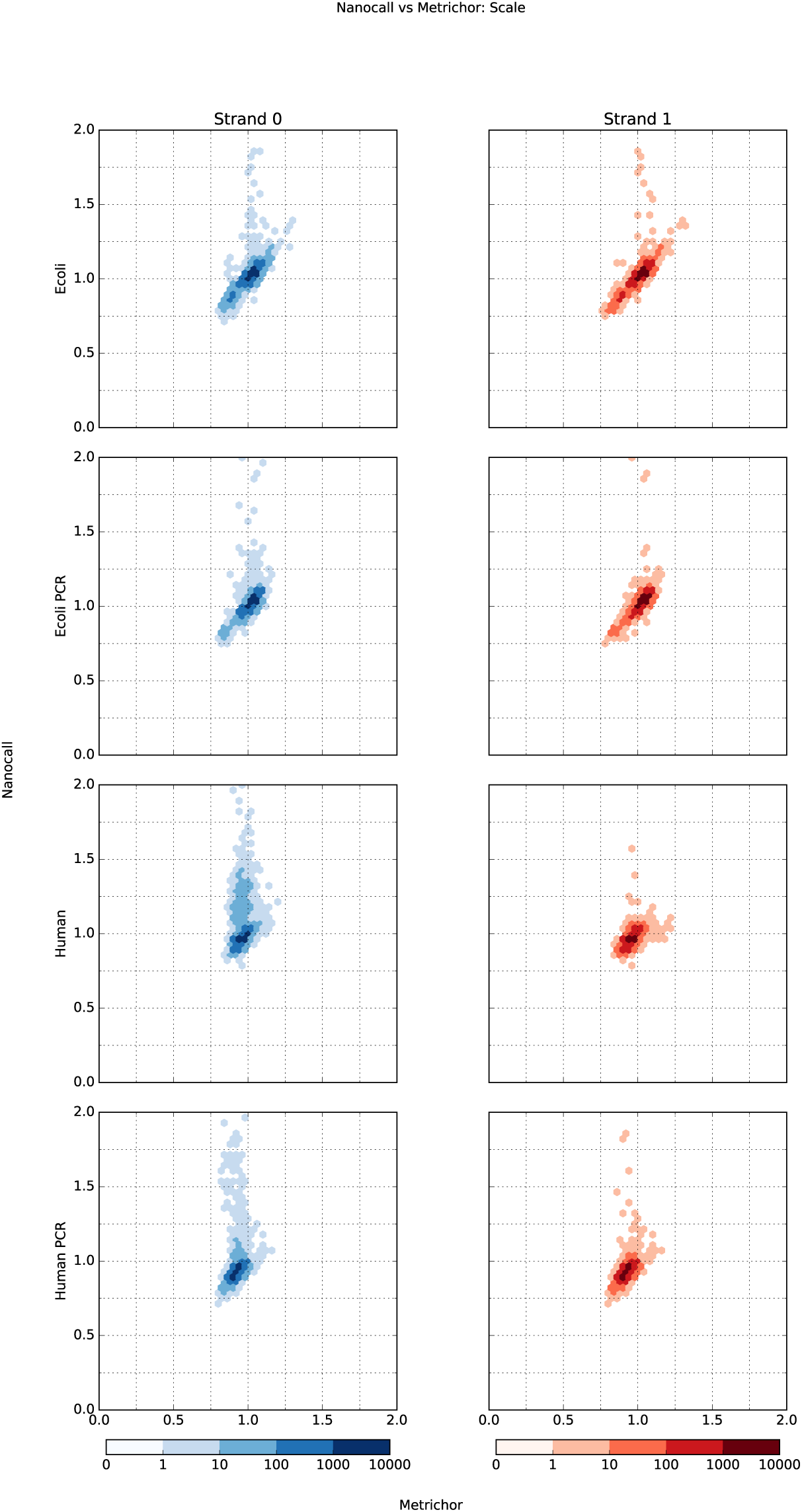
Nanocall vs Metrichor `scale`.

**Fig. 2.**
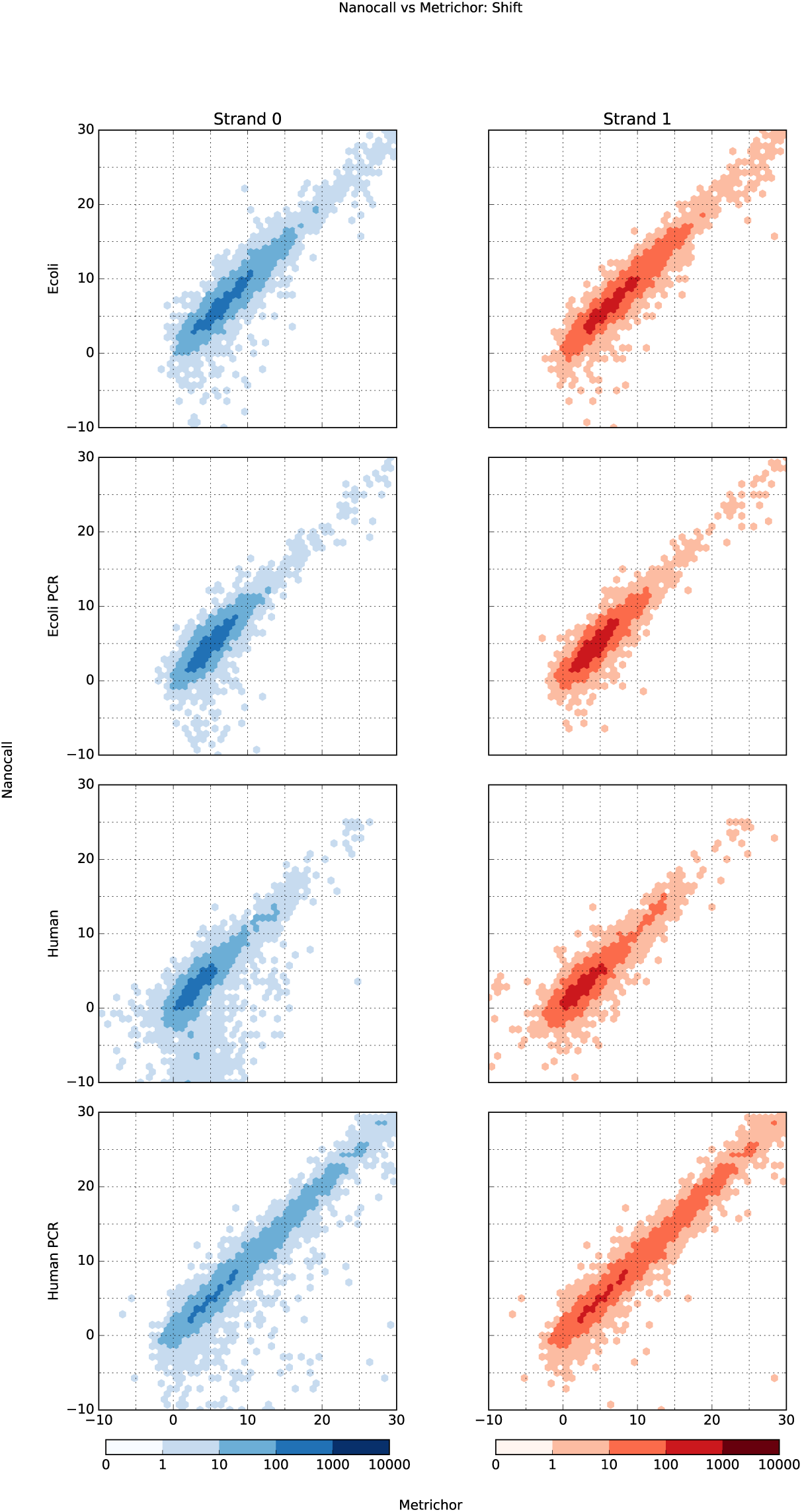
Nanocall vs Metrichor shift.

**Fig. 3.**
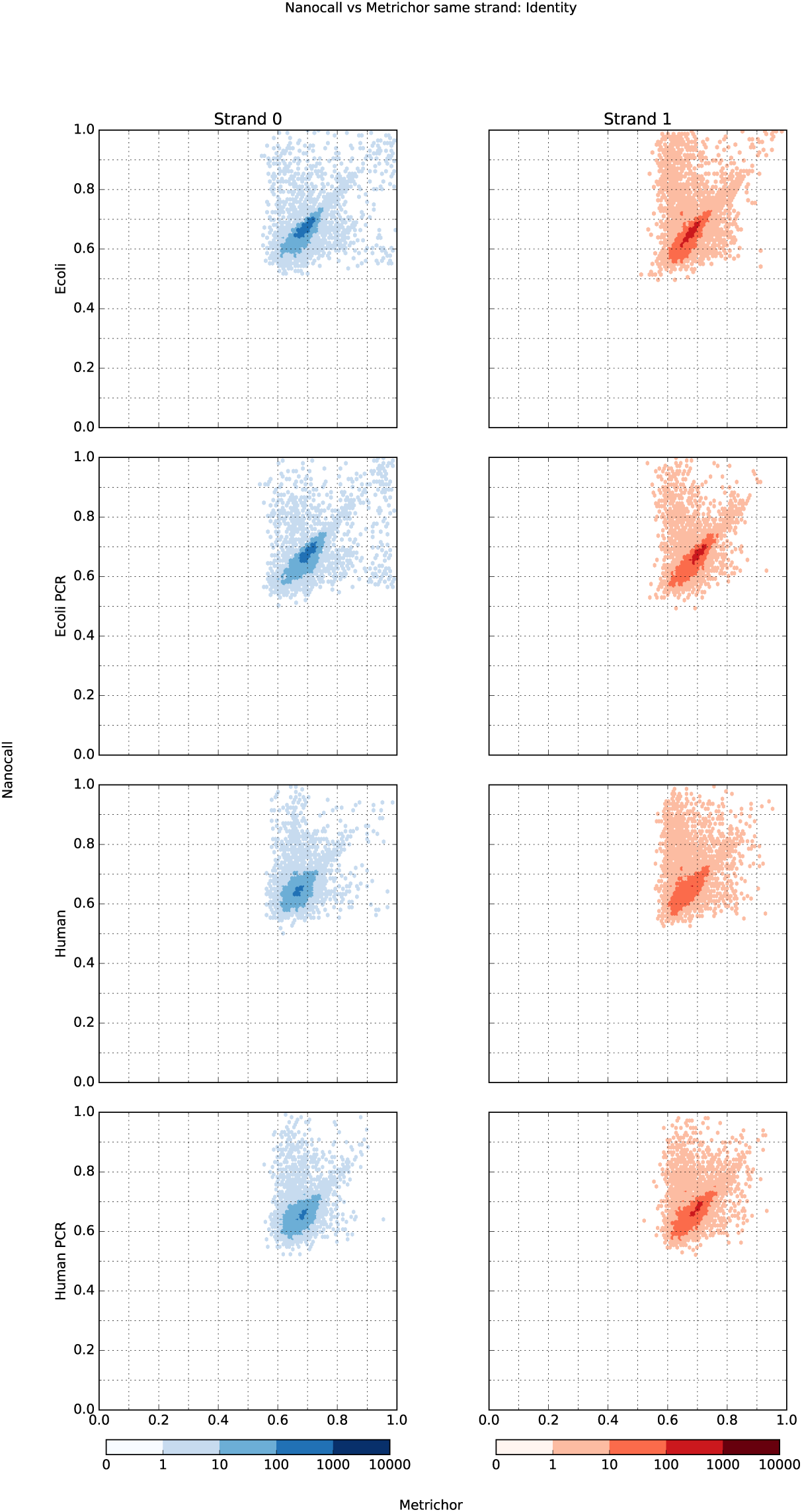
Nanocall vs Metrichor identity.

**Fig. 4.**
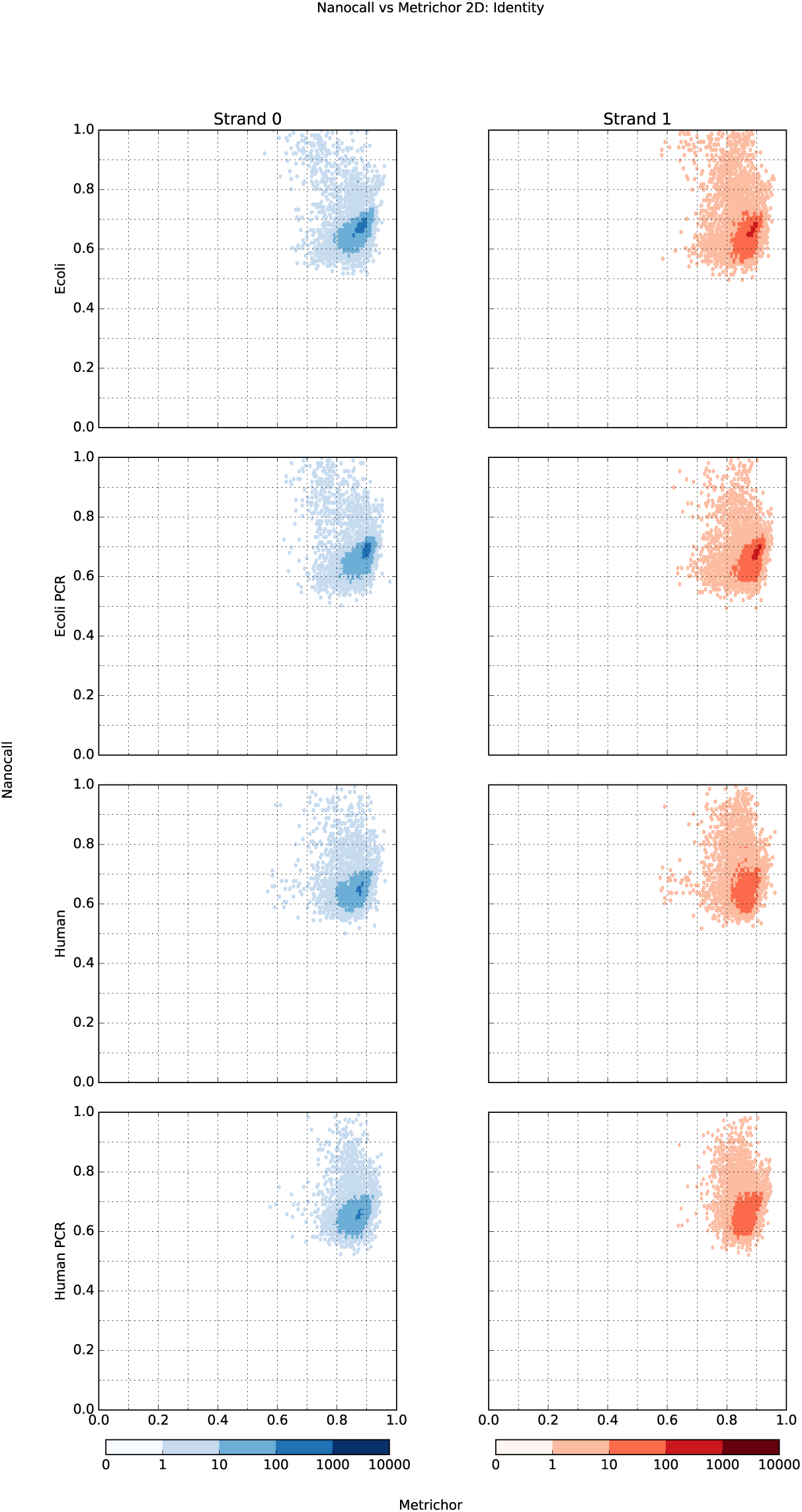
Nanocall vs Metrichor ID identity.

**Fig. 5.**
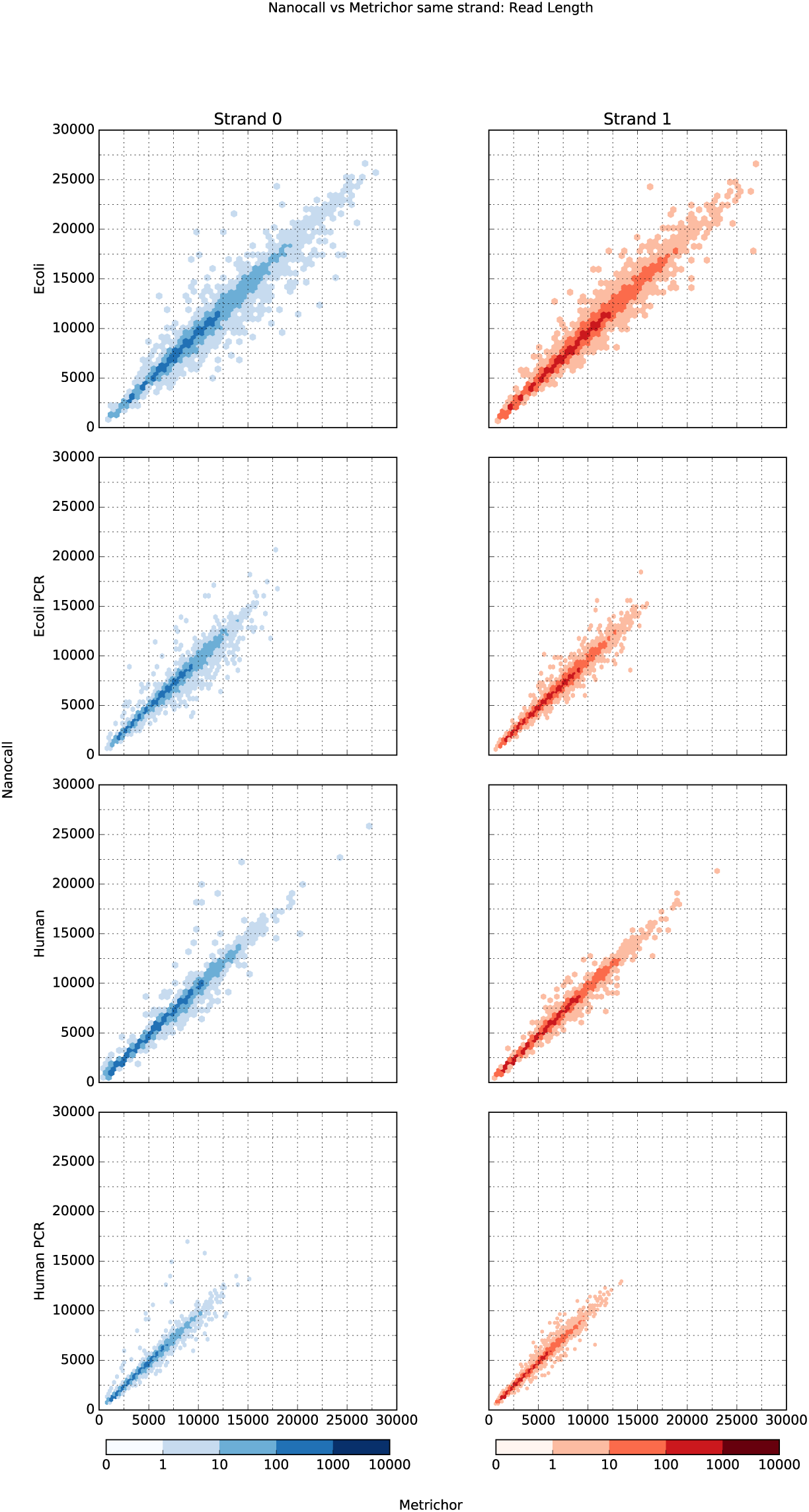
Nanocall vs Metrichor read length.

**Fig. 6.**
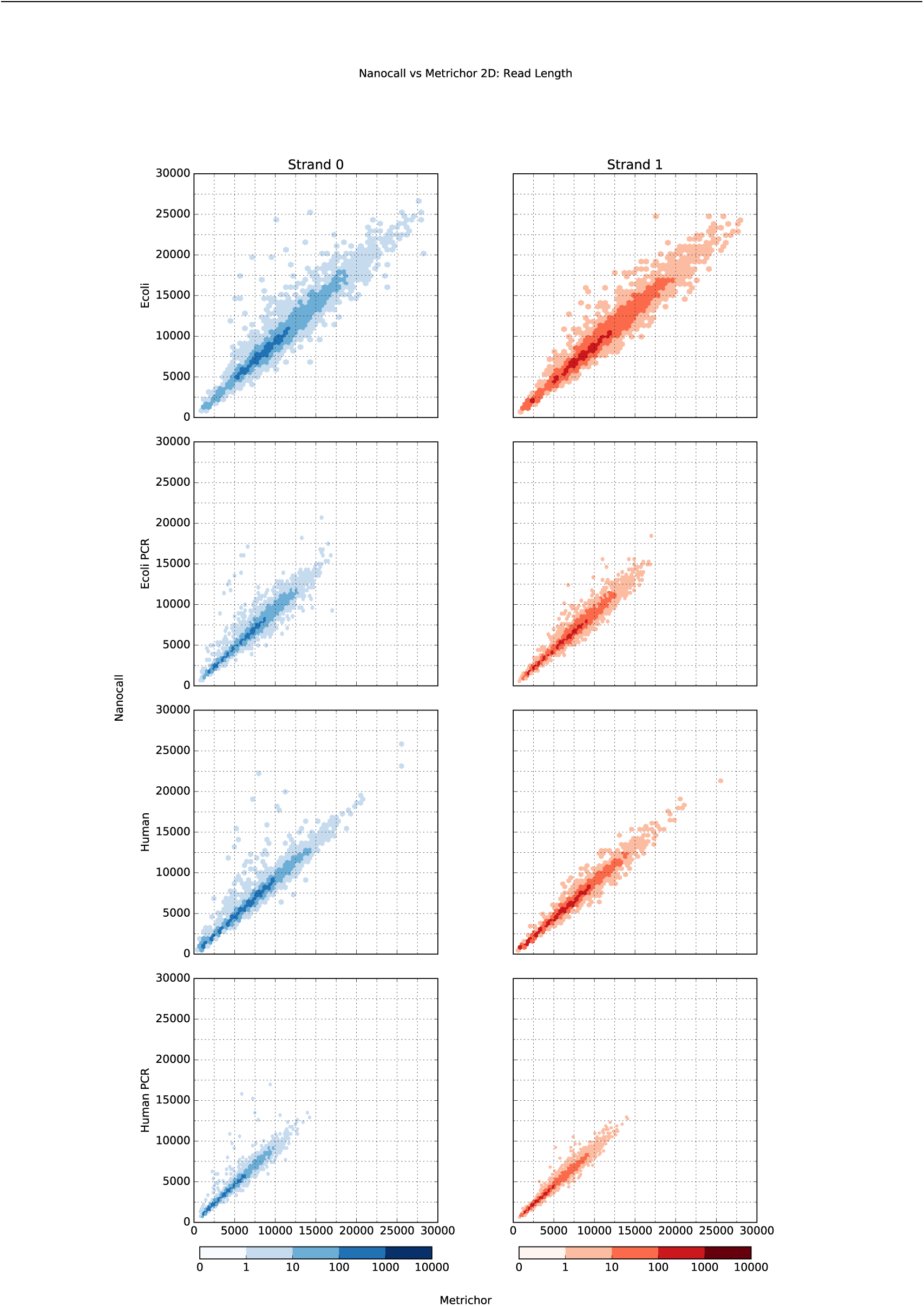
Nanocall vs Metrichor 2D read length.

**Fig. 7.**
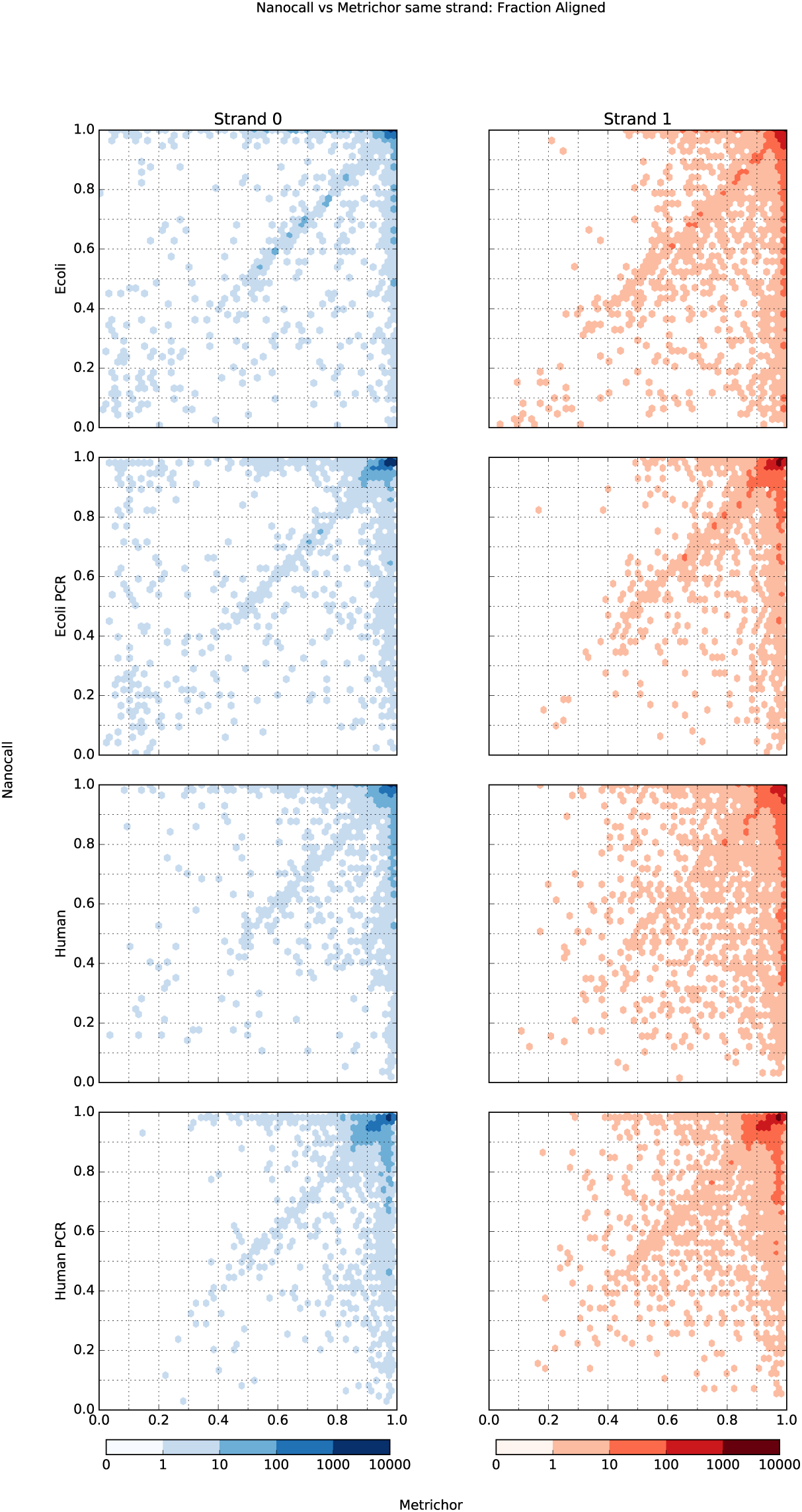
Nanocall vs Metrichor fraction aligned.

**Fig. 8.**
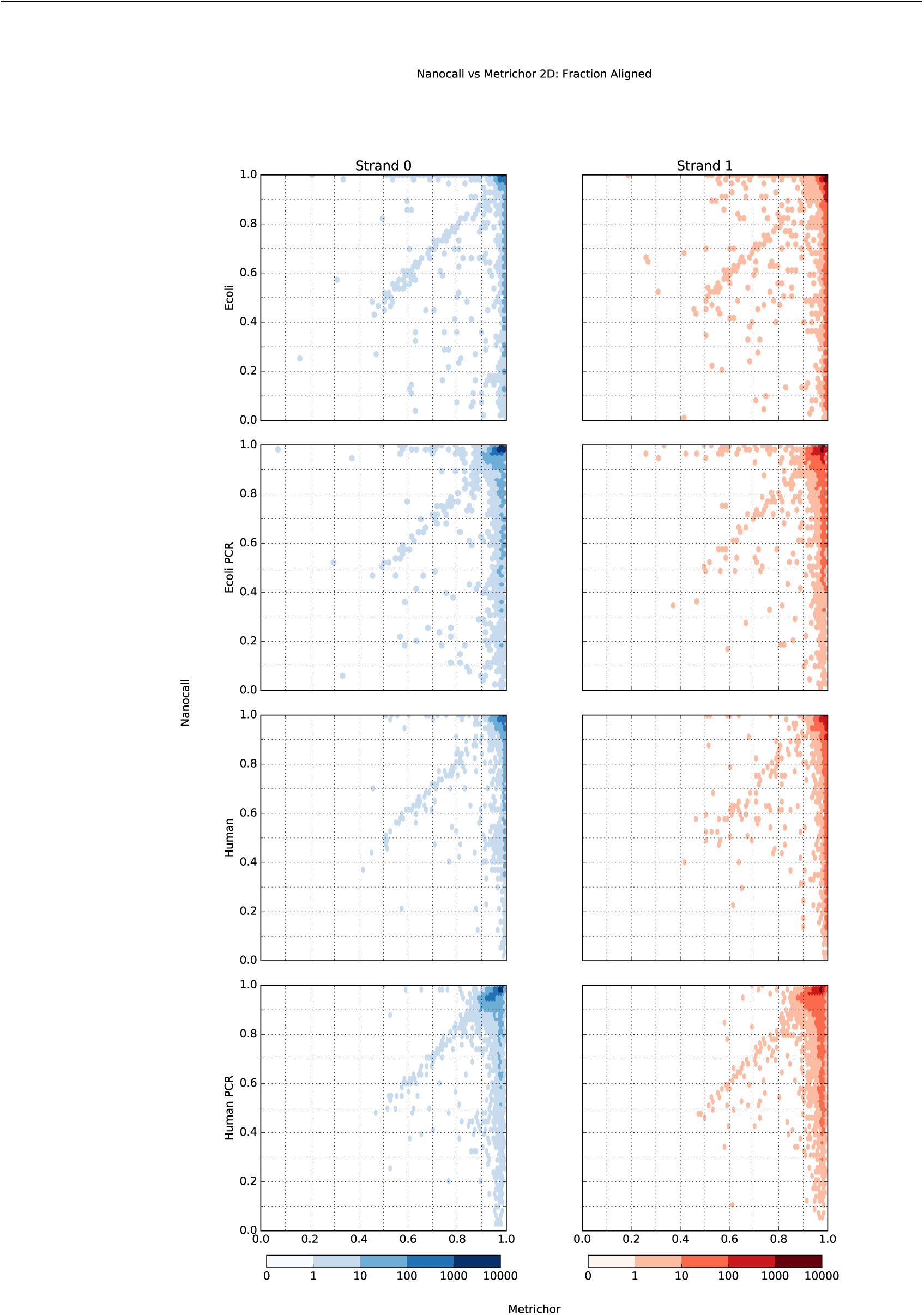
Nanocall vs Metrichor 2D fraction aligned.

**Fig. 9.**
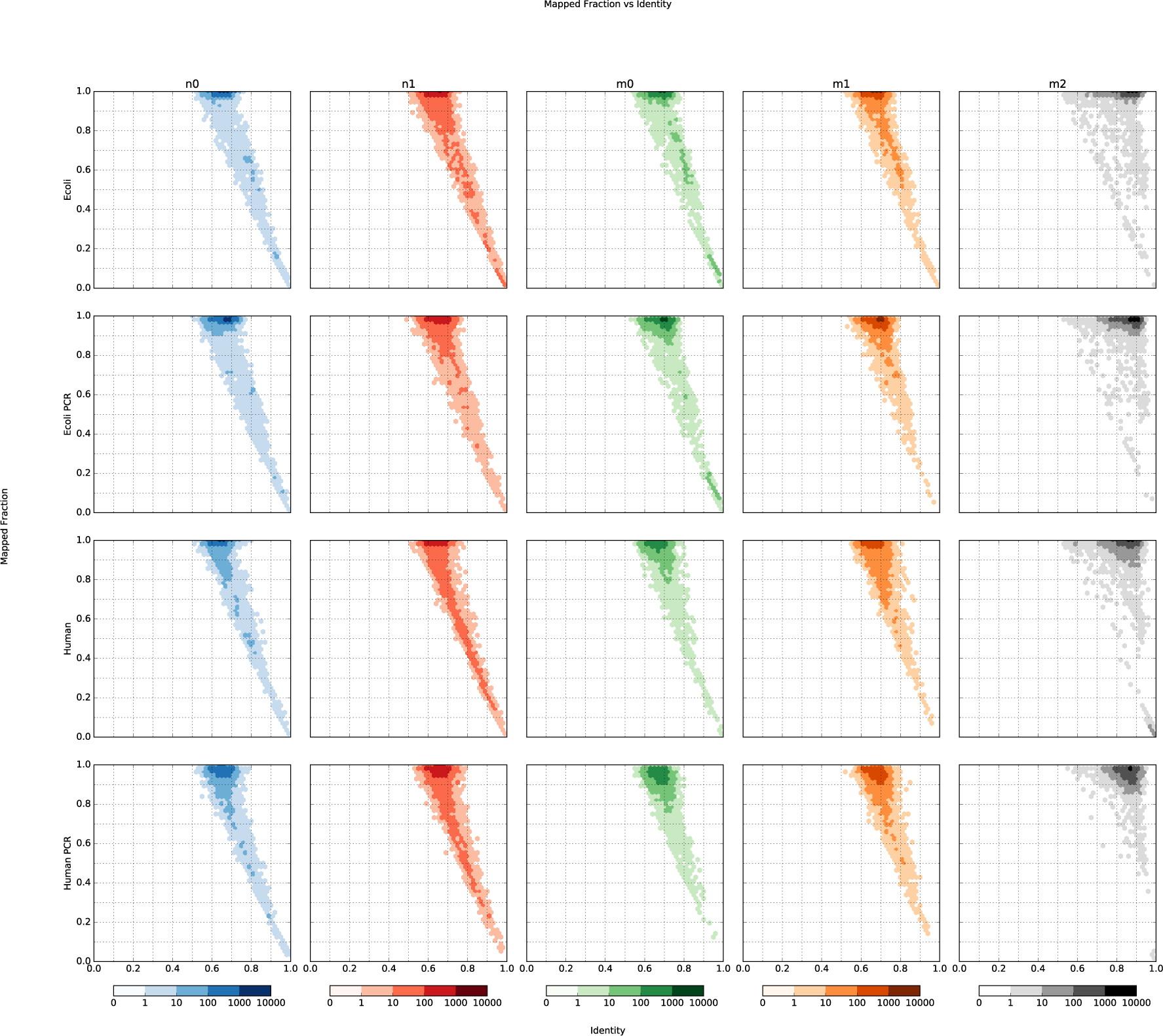
Identity vs fraction aligned, for each of the 5 read types: Nanocall template/complement, Metrichor template/complement/2D.

To quantify the effects of the default transition parameters, we used 1,000 human PCR reads, and ran Nanocall in double strand scaling mode without transition parameter training (`2ss-no_tt`), using several parameter values: *p*_stay_ ∈ {.9,1.0*, 1.1} and *p*_skip_ ∈ {2.6, 2.8, 3.0*, 3.2} (* denotes the default). The results of these runs are given in Table 2, showing that, while *p*_stay_ and *p*_skip_ do affect mappability, their influence is quite limited: a difference of < 1% in mappability between all runs. Note: we expect *p*_stay_ and *p*_skip_ to have negligible effect when transition parameter training is enabled (2ss).

**Table 2.**
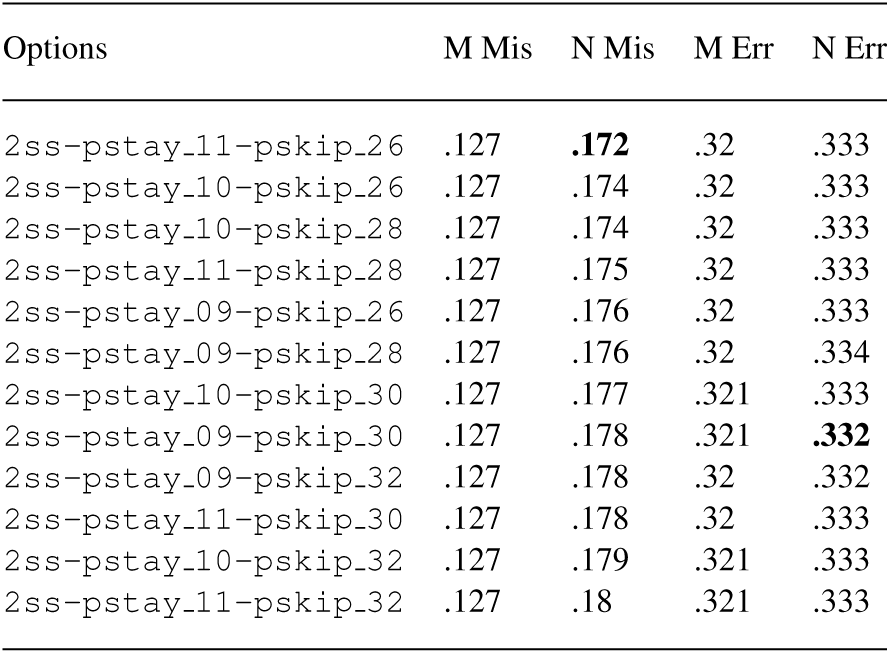
Influence of default transition parameters *p*_stay_ and *p*_skip_. All runs on 1,000 human pcr reads, with double strand scaling with no transition parameter training (`2ss-no_tt`). `pstay_xy`: *p*_stay_ = *x.y*; `pskip_uv`: *p*_skip_ = .*uv*. Column labels: see Table 1.

We also studied the effects of perturbing the parameters that control training: the number of events used (default 100), the maximum number of rounds per strand (default: 10), and the minimum improvement in fit (default: additive term of 1.0 in log space, corresponding to multiplicative term *e*). The results are given in Table 3. Clearly, increasing the number of events used per strand (`nume`) directly improves mappability, but also decreases speed. Increasing the maximum number of training rounds (`maxr`) has the same effect, but improvements in mappability are lost beyond 10 rounds. The effect of the minimum fit improvement (`minp`) on mappability and even speed is harder to quantify.

**Table 3.** Influence of parameters that control training. Runs on 1,000 human PCR reads, using double strand scaling mode with transition parameter training (`2ss`). `nume_x`: use *x* events used per strand; `maxr_y`: maximum of *y* rounds per strand; `minp_uv`: minimum fit improvement of *u.v* in log space. Column labels: see Table 1.

| Options | M Mis | N Mis | M Err | N Err | Speed |
| --- | --- | --- | --- | --- | --- |
| 2ss-nume_250 | .127 | **.165** | .32 | .331 | 93 |
| 2ss-minp_15 | .127 | .175 | .32 | .332 | 137 |
| 2ss-maxr_10 | .127 | .177 | .321 | .333 | 85 |
| 2ss-minp_10 | .127 | .177 | .321 | .333 | 113 |
| 2ss-nume_200 | .127 | .177 | .321 | .333 | 121 |
| 2ss-maxr_05 | .127 | .178 | .321 | .333 | 28 |
| 2ss-maxr_15 | .127 | .178 | .321 | .333 | 46 |
| 2ss-maxr_20 | .127 | .178 | .321 | .333 | 46 |
| 2ss-minp_20 | .127 | .181 | .32 | .333 | 147 |
| 2ss-maxr_04 | .127 | .184 | .32 | .332 | 43 |
| 2ss-minp_05 | .127 | .189 | .32 | .332 | 34 |
| 2ss-nume_150 | .127 | .191 | .32 | **.33** | 72 |
| 2ss-maxr_03 | .127 | .2 | .32 | .333 | 137 |
| 2ss-maxr_02 | .127 | .221 | .32 | .337 | 76 |
| 2ss-nume_100 | .127 | .221 | .32 | .331 | 223 |
| 2ss-nume_050 | .127 | .27 | .319 | .332 | **370** |

## 5. CONCLUSION

In this work we presented Nanocall, an open source, MIT licensed basecaller for data produced by Oxford Nanopore MinION instruments. Nanocall uses some simple heuristics for splitting the sequence of events (current levels) into strands, it models the events using a hidden Markov model where the states are the kmers being sequenced, it optionally scales the pore model emissions using several rounds of Expectation Maximization based on posteriors computed with Forward-Backward, and it produces basecalls by running Viterbi.

Overall, Nanocall produces reads comparable in mappability and quality to Metrichor 1D reads with ~ 68% identity. As an important technical difference from Metrichor, with Nanocall we found that double-strand pore model scaling seems to work better than single-strand scaling, suggesting that the latter might lead to model overfitting.

## ACKNOWLEDGEMENTS

We thank Tim Massingham from Oxford Nanopore Technologies for describing how Metrichor’s basecaller trains the scaling parameters. We used this document as a starting point for the derivation of our training method provided in the supplement.

